# Chronic morphine treatment leads to a global DNA hypomethylation via active and passive demethylation mechanisms in mESCs

**DOI:** 10.1101/2025.03.16.643512

**Authors:** Manu Araolaza, Iraia Muñoa-Hoyos, Itziar Urizar-Arenaza, Irune Calzado, Nerea Subirán

## Abstract

Epigenetic changes are essential for normal development and ageing, but there is still limited understanding of how environmental factors can cause epigenetic changes that leads to health problems or diseases. Morphine is known to pass through the placental barrier and impact normal embryo development by affecting the neural tube, frontal cortex and spinal cord development, and, as a consequence, delaying nervous system development. In fact, in-utero morphine exposure has shown alterations in anxiety-like behaviours, analgesic tolerance, synaptic plasticity and the neuronal structure of offspring. However, how morphine leads to abnormal neurogenesis and other physiological consequences during embryo development is still unknown. Considering that DNA methylation is a key epigenetic factor crucial for embryo development, our aim is to elucidate the role of methylation in response to morphine. Chronic morphine treatment (24h, 10μM) induces a global hypomethylation in mESC. WGBSeq identifies 16,808 sensitive to morphine which are involved in embryo development, signalling pathways, metabolism and/or gene expression, suggesting that morphine might impact methylation levels at developmental genes. Integrative analyses between WGBSeq and RNASeq identified *Tet1* as morphine-sensitive gene. Morphine increased the gene expression of *Tet1*, modifying the methylation levels at the promoter. On the other hand, RNASeq and qRT-PCR analyses revealed that *Dnmt1* gene expression decreased after morphine treatment, without altering the methylation patter at its promoters. By MS/MS approaches confirms a decrease in DNA methylation after chronic morphine treatment, together with an increase in hydroxymethylation global levels in mESCs. In conclusion, morphine induces a global hypomethylation in mESC through different mechanisms that involves passive demethylation and a self-regulatory mechanism via active demethylation.

**One Sentence Summary:** Understanding how environmental epigenetics impact on embryo development aids to unravel changes that leads to health problems or diseases in the adulthood.

**Graphical abstract:** Chronic morphine treatment induces a global hypomethylation in mESC through different mechanisms that involves passive demethylation and a self-regulatory mechanism via active demethylation in genes involved in embryo development.

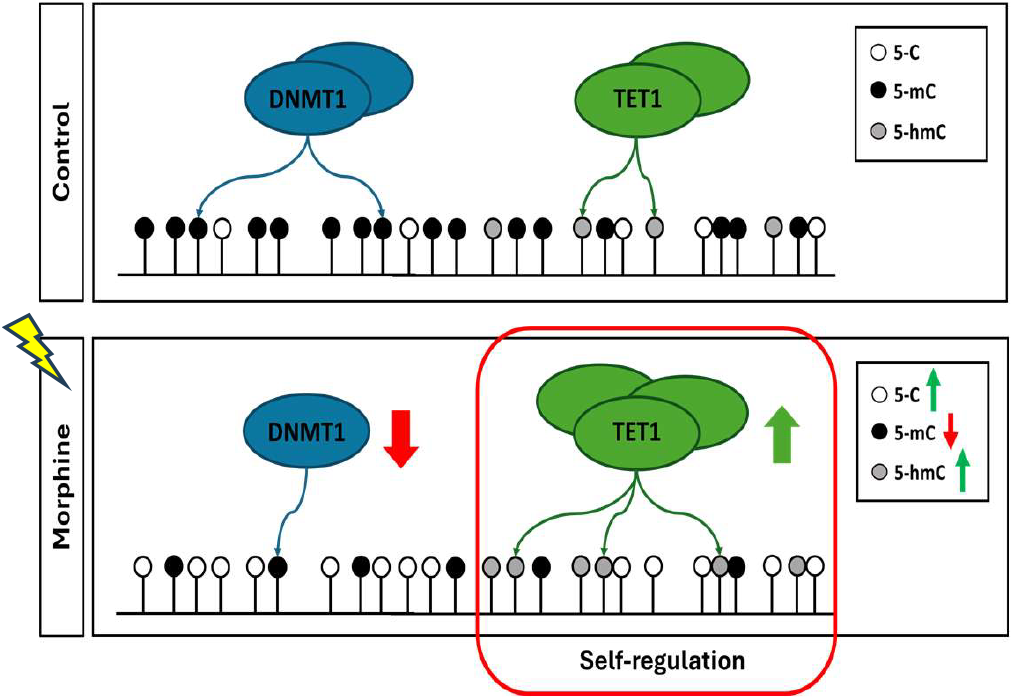

## INTRODUCTION

Nowadays, there is a greater understanding of how various environmental factors affect our lives and health. However, the underlying mechanisms involved remain largely unknown. While epigenetic changes are crucial for development and aging, some of these changes can lead to health issues or diseases, including autoimmune disorders, neurodevelopmental syndromes, cardiovascular conditions, and cancer (de Gonzalo-Calvo et al., 2017; Moosavi & Motevalizadeh Ardekani, 2016). Notably, prenatal developmental processes are highly sensitive to toxic chemicals and stress, suggesting that environmental factors can disrupt embryo development and result in organ dysfunction after birth (Ko et al., 2019). Morphine, an addictive drug, is widely used in modern medical practice. Its primary therapeutic value lies in its ability to provide pain relief, or analgesia(D Matthes et al., 1996; Manglik et al., 2012). However, it is well-documented that morphine has numerous adverse physiological side effects (Nakatani, 2017). Morphine can readily cross the placental barrier and, as a result, reach the embryo (Kazemi et al., 2011; Levitt, 1998), leading to various problems during the developmental process. Morphine has been reported to reduce the weight of several organs during development—such as the brain, kidneys, and liver in rats (Eriksson & Rönnbäck, 1989), but the most significant are associated with disruptions in normal neural development, causing delays in nervous system maturation (Sahraei et al., 2013). Furthermore, in utero exposure to morphine has been linked to alterations in anxiety-like behaviours, analgesic tolerance, synaptic plasticity, and the neuronal structure of offspring (Byrnes et al., 2011; Gapp et al., 2014). Although it is well established that morphine’s effects are mediated through opioid receptors (Subirán et al., 2011), the precise mechanisms by which morphine impacts embryonic development remain unclear.

DNA methylation confers marks of transcriptionally silent chromatin at developmental genes during embryo development, as it is the mechanism that defines the direction of differentiation that each cell (Correa et al., 2020; Smith & Meissner, 2013). This path is defined during the process of embryogenesis, where the stem cells ensure their renewal (Toyooka, 2021). The maintenance of the methylation pattern of DMR sites is a prerequisite for normal mammalian development, and their loss causes serious damage to the developmental process in mammals (Howell et al., 1998, 2001; McGraw et al., 2013). For example, mice embryos lacking the DNA methyltransferases DNMT1 (*Dnmt1-/-)* die, as development process is disrupted caused by a deregulation in DNA methylation pattern (E. Li et al., 1992). Although DNA methylation is considered a long-lasting epigenetic mark, it can undergo gradual passive loss if the methylation pattern is not maintained during DNA replication across cell generations (Inoue et al., 2011). Therefore, passive DNA does not require specific protein machinery, but DNA demethylation can also occur through an active process mediated by the ten-eleven translocation (TET) protein family (Wu & Zhang, 2014; Zhao & Chen, 2013). DNA demethylation is also a key mechanism in embryogenesis, but in this case, to ensure the pluripotency of cells (Bagci & Fisher, 2013). Zygotes resulting from the fertilization of gametes undergo sequential divisions to drive pluripotent stem cells (PSCs) in the early stages of the embryo, in which DNA demethylation is crucial to ensure the pluripotency of those cells (Bagci & Fisher, 2013). Considering that DNA methylation pattern is a key factor crucial for embryo development, our aim is to elucidate the role of this epigenetic mark and its regulatory machinery in gene expression in response to morphine.

## RESULTS

### Effect of chronic morphine treatment on DNA methylation in mESCs by *Whole Genome Bisulphite Sequencing* (WGBS)

First, we want to determine the methylation changes induced by chronic morphine treatment in mESCs *in vitro*. mESCs expressing GFP protein under the Oct4 promoter were treated with morphine treatment at 10mM for 24 h (Yang et al., 2019), which did not induce any morphological changes on mESCs (**Fig. 1A**). We next studied in depth the distribution of methylated DNA throughout the whole genome by WGBS. Considering the differentially enriched regions with p < 0.05 and FDR < 0.05, a total of 203,337 and 223,280 Differentially Methylated Cytosines (DMCs) were annotated by using two different statistical tools, *edgeR* and *methylkit* respectively. Through the integration of these acquired data, 153,352 DMCs common to both tools were identified, representing 56.12% of the total DMCs that were identified. (**Fig. 1B**). To increase the biological relevance of the changes, we considered a fold change of 2 in *edgeR* data, and 20% percent-difference changes in *methylkit* data, reducing up to 78,235 common. Remarkably, mainly hypomethylated cytosines were identified, which represent the 72.1% of the common DMCs, meanwhile hypermethylated cytosines only reached the 27.9% of them. Thus, morphine led to a decrease on whole genome DNA methylated regions (**Fig. 1B**). Because CpG islands (CGIs) have been implicated in methylation dependant gene expression regulation (Cain et al., n.d.; Deaton & Bird, 2011; Illingworth & Bird, 2009), we also analysed the changes induced by morphine at CGIs and flanking features. More than 90% of DMCs were found in open sea area, but CpG islands and surrounding areas were remarkably enriched with hypermethylations. Although we observed an overall decrease of DNA methylation, hypermethylated DMCs were specifically reported at CGIs and shore regions after morphine treatment (**Fig. 1C**). As DNA methylation distribution at promoters also impact on gene expression (Weber et al., 2007), we also analyzed the distribution of these differentially methylated cytosines at those regions. Remarkably, we observed a greater presence of hypermethylated DMC at promoter regions than other region, a process that may have a very large relevance in the transcription levels of the associated genes (**Supp. fig. 1**).

**Figure 1.**
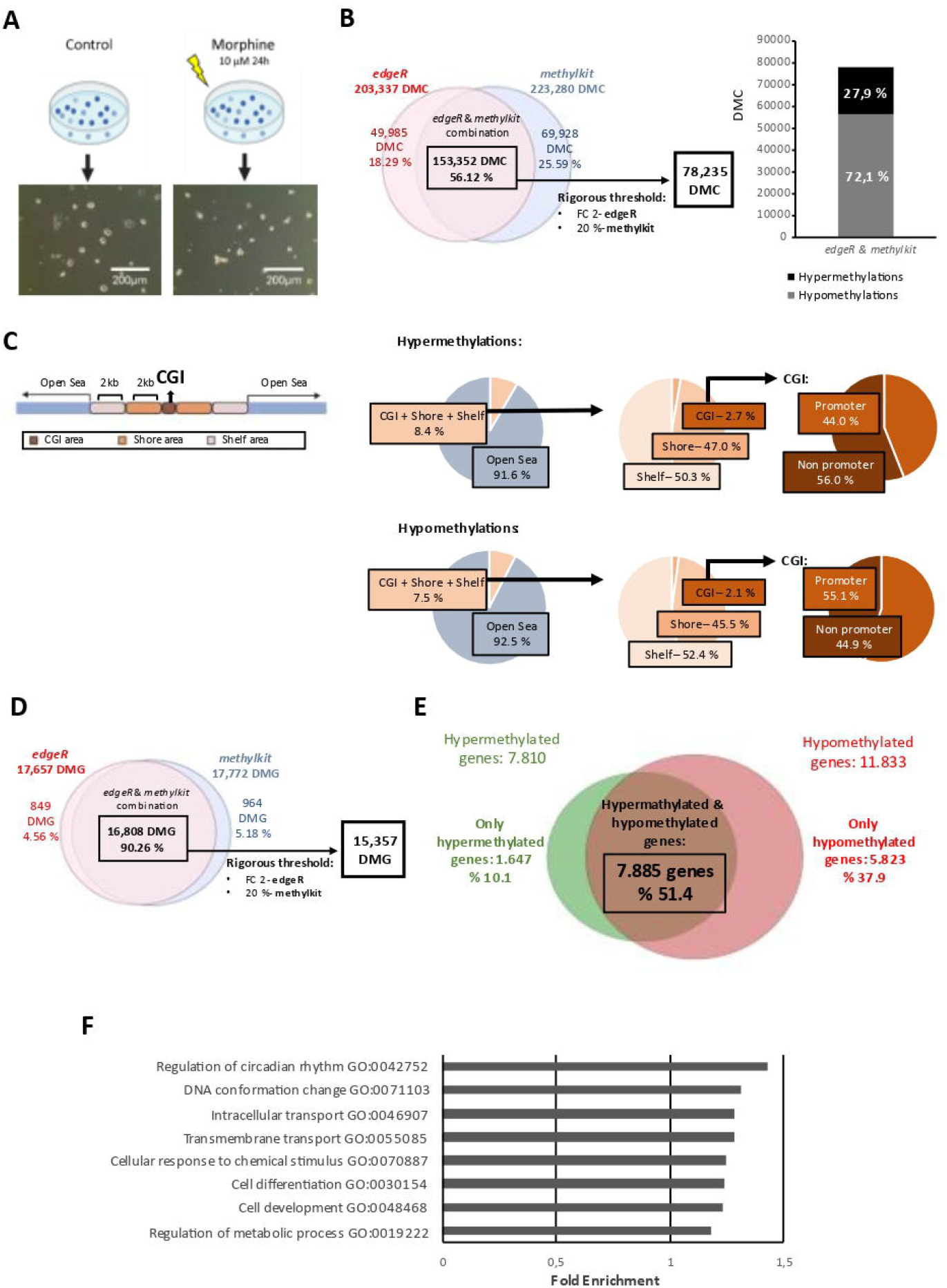
DNA methylation distribution after chronic morphine treatment in mESC by using WGBS. **(A)** Schematic representation of mESCs culture and 24 h morphine treatment for *in vitro* epigenetic changes detennination. Representative image of treated and untreated mESCs is also shown. Scale bar = 200µm. (B) *Differentially methylated cytosines* (DMC) identified through edgeR and methylKit tools after chronic morphine treatment. Venn diagram show data integration between both tools. Percentages of hype1methylations and hypomethylations of DMCs fo1m integrative data. **(C)** Pie-chart showing CpG feature distribution of DMC in CpG island after chronic morphine treatment at CGis regions (belonging to promoter or non-promoter region+ **1** kb from TSS), shore (< 2 kb), shelf (< 4 kb) and open sea (the rest of the genome). **(D)** Vem1 diagram representation of *differentially methylated genes* (DMG) identified from both *edgeR* and *methylKit* tools and **(E)** which of in common DMG are exclusively hype1methylated and hypomethylated genes or show both hypennethylations and hypomethylations. **(F)** Functional emichment analysis showing the most indicative biological functions of the DMG genes. Statistical analyses Fisher’s analyses forp >0.05.

Next, we evaluated how many of these DMCs fall exactly in genes to define the Differentially Methylated Genes (DMGs) induced by morphine. This allowed us to gain a comprehensive view and identify all Differentially Methylated Genes (DMGs). According to our data, *edgeR* identified 17,657 DMGs, while *methylKit* identified 17,772 DMGs. To ensure the reliability of our results, we decided to focus exclusively on the 16,808 DMGs that were consistently identified by both tools, representing a 90.26% overlap (**Fig. 1D**). To further enhance the relevance of the data, we applied more stringent thresholds: a fold change of 2 in *edgeR* and a 20% percent-difference threshold in *methylKit*. As a result, the dataset was refined from 16,808 DMGs to 15,357 DMGs (**Fig. 1D**). Chronic morphine treatment predominantly induced hypomethylation within genes, a pattern consistent with the observations for genome-wide DMCs. Most affected genes (51.4%) exhibited both hypermethylation and hypomethylation. However, in 37.9% of cases hypomethylations were exclusively observed in genes related to basic cellular processes such as signalling transduction, apoptosis, metabolism as well as developmental processes, while only 10.1% of the genes were exclusively hypermethylated, exclusively involved in primary metabolic processes (**Fig. 1E, Supp fig 2**). To understand the biological functions in which morphine was involved, *Gene Ontology* (GO) analysis was performed. Functional enrichment analysis showed that genes sensitive to morphine were involved mainly in regulation of circadian rhythm, DNA conformation change, cell differentiation and development and metabolic processes (**Fig. 1F**).

**Figure 2.**
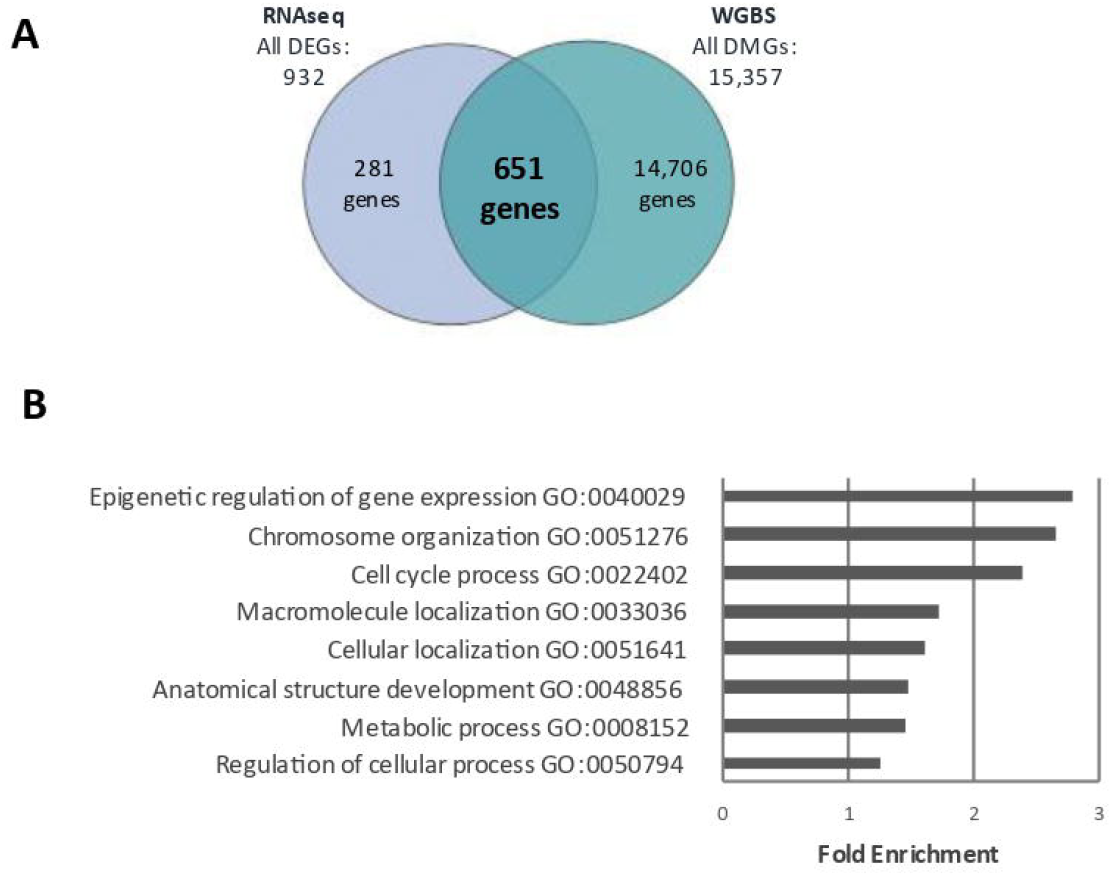
Morphine-sensitive genes from integrntive analyses ofRNAseq and WGBS data. **(A)** Venn diagram representation showing the overlap between *differentialy methylated genes (DMG)* fonn WGBS data, and RNA-seq identified *differentially expressed genes* DEGs after chronic morphine treatment. (B) Gene Ontology analysis showing the top biological functions, perfo1med with the criteria of Fisher’s method for *p<0*.*05*.

Aiming to understand the relevance of methylome on transcriptomic deregulation after chronic morphine treatment, an integrative analysis was performed between the WGBS and RNA-Seq data. For that purpose, we used RNA-Seq data generated previously by our research group, in which we described the global gene expression profiling after chronic morphine treatment (GEO Store: GSE151234) (Muñoa-Hoyos et al., 2020). Briefly, a total of 932 differentially expressed genes (DEGs) were identified after 24 h of morphine treatment (**Fig. 2A**), mainly involved in nuclear and cell division, DNA repair, chromosome organization, gene expression, metabolism and signaling process, among others (Muñoa-Hoyos et al., 2020). Venny diagram identified 651 genes in common between both DEGs and DMGs, which were involved in epigenetic regulation, chromosome organization, cell cycle process, among others (**Fig. 2B**). Specifically, 24 genes involved in epigenetic regulation and gene expression were identified, including core genes belonging to DNA methylation machinery (**Table 1**). Remarkably, DNA (cytosine-5)-methyltransferase 3-like (*Dnmt3l*) as well as methylcytosine dioxygenases (*Tet1*) were identified as a morphine-sensitive genes as transcriptomic and methylome levels, which may indicate morphine can potentially autoregulate DNA methylation levels.

**Table 1.**
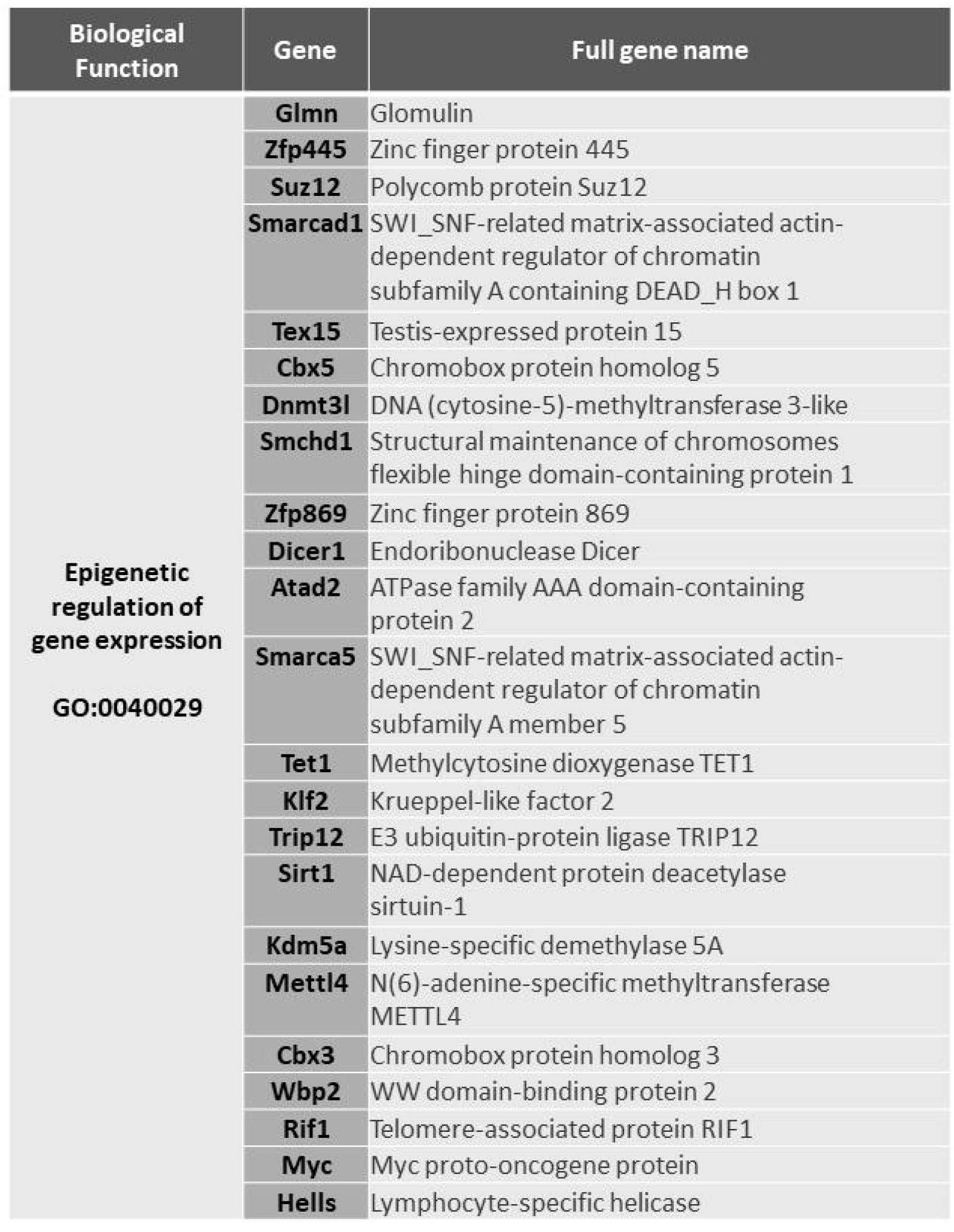
Morphine-sensitive genes related to biological function *Epigenetic regulation of gene expression (GO:0040029)* that have been identified by integrating RNAseq and WGBS databases.

### Effect of chronic morphine treatment on DNA methylation machinery

WGBS and RNA-Seq approaches confirmed that morphine was able to regulate the DNA methylation machinery in mESCs (**Fig. 3, Supp fig 2, 3**). The levels and patterns of DNA methylation are regulated by both DNA methyltransferases (DNMT1, DNMT3A and DNMT3B) and demethylating proteins, including the ten-eleven translocation (TET) family of dioxygenases (TET1, TET2 and TET3) (Zeng & Chen, 2019). Landscapes of those genes obtained from the UCSC genome browser showed an effect of chronic morphine treatment not only on Tet1 and Dnmt3l but also on Dnmt1 gene expression (Fig. 3 Supp fig 4A). In regard to ‘demethylating’ enzymes, morphine led to a decrease in DNA methylation level at the Tet1 gene that was consistent with its an up-regulated expression. Morphine mainly induced hypomethylation along the whole *Tet1* gene, including the in two promoters of the gene (**Fig 3. A, C**). Consistent with the decrease on DNA methylation distribution, we observed an upregulation of the gene that involved both *Tet1* transcripts (**Fig 3. A, C**). WGBS also proved morphine altered DNA methylation pattern on *Tet3* at different regions, including at promoter areas, which did not impact on its gene expression. Finally, chronic treatment did neither impact on *Tet2* genes at both methylation nor gene expression levels (**Supp Fig. 3**).

**Figure 3.**
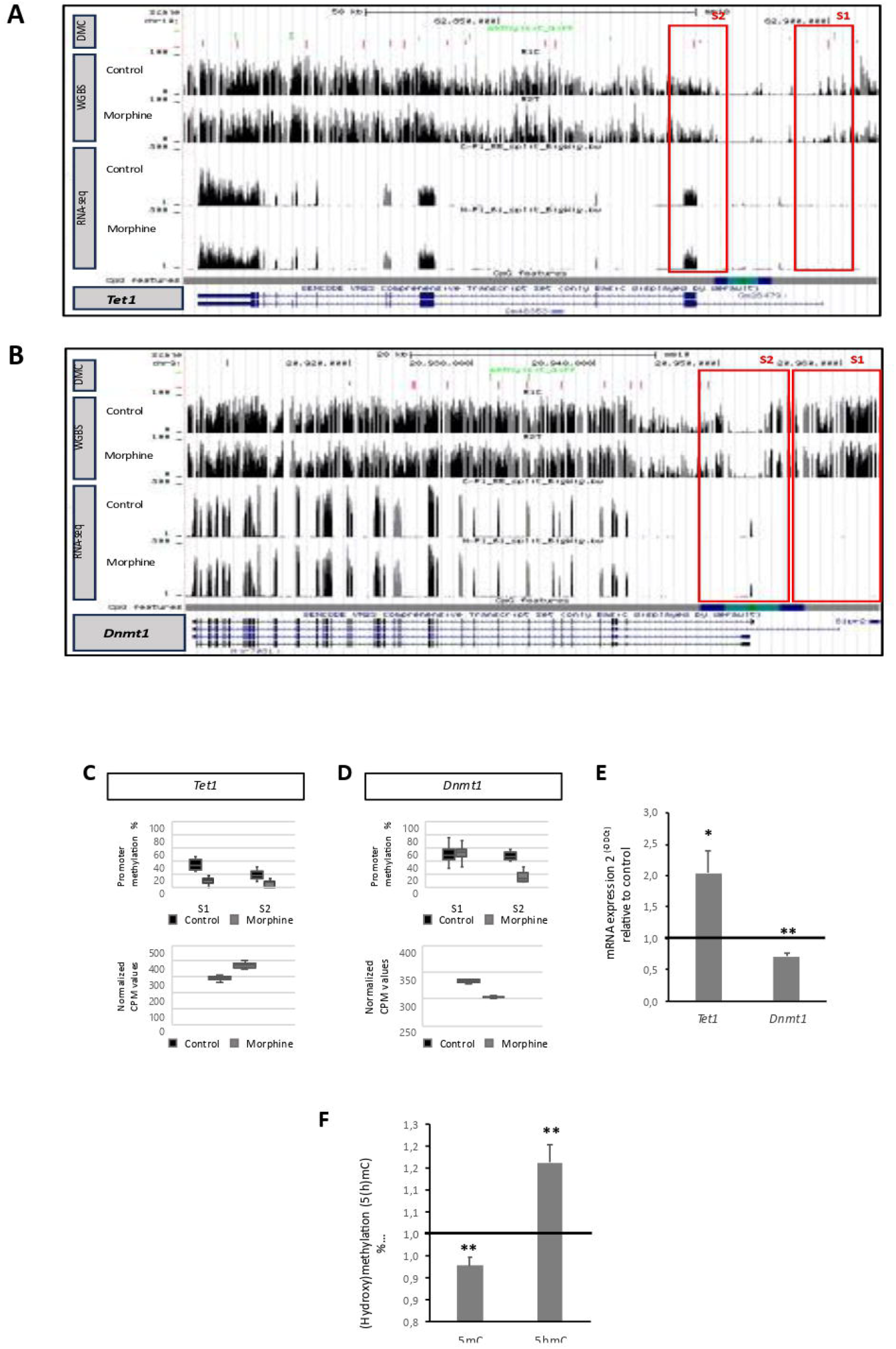
Effect of chronic morphine treatment on DNA methylation/demethylation machinery. RNA-seq and WGBS track for **(A)** DNA demethylating enzyme *Teti* gene and **(B)** DNA methyltransferase *Dnmti* gene. CpG features track was composed by CpG islands in green, shores in light blue, shelfs in dark blue and open sea in grey. Red boxes point out the enrichment and gene expression change at promoters. Box and whisker plot showing the percentage of methylation at promoters and CPM values for **(C)** *Teti* gene and **(D)** *Dnrnti* gene after chronic morphine treatment. **(E)** RT-qPCR analysis for the validation of *Teti and Dnmti* gene expression. *Gapdh* and *Pcx* were used as housekeeping genes for mRNA level analysis and acquired Ct values were nonnalized respect to the control sample using 2^ddcr^ (n=5). (F) DNA methylation and hydroxymethylation levels after chronic morphine treatment by MS/MS. Percentage of 5mC and 5hmC levels nonnalized with respect to control (n=5). Statistical analysis was performed using the Student’s t-test, differentiating the level of significance: *p<0.05, **p<0.01 and ***p<0.001.

Morphine also modified gene expression and DNA methylation level on Dmt3l gene (**Suppl fig 4. A, D**). Similar to observed in *Tet1* gene, morphine induced an up regulation of *Dmt3l* gene expression, which is consistent with a downregulation of DNA methylation at promotor (S3) in charge of the majority of transcripts that are expressing. Athough DNMT3L protein is part of the DNA methylation machinery, it does not directly involved on catalytic activity (Guenatri et al., 2013; Kareta et al., 2006). Therefore, we next evaluated therefore, the morphine impact on DNA methyltransferases. Contrary with the results obtained for TET proteins, morphine decreased DNA methylation levels of *Dnmt1* along the whole gene, including at promoter regions, but this alteration was no consistent with its reduced expression (**Fig 3. B, D**). Specifically, *Dnmt1* landscape showed hypomethylations induced by morphine at one of the promoters (S2; principal promoter area), which is in charge of expressing 3 out of 4 *Dnmt1* transcripts and correspond to a GGI region. No methylation changes were observed on the other promotor region of the gen upstream (S1). The landscape of the RNA-Seq, however, showed a downregulation of the *Dnmt1* gene expression, which did not correspond to the DNA methylation pattern induced by morphine (**Fig 3. B, D**). Whit respect to other DNA methyltransferases, morphine mainly induced hypermethylation along the whole *Dnmt3a* gene, which are not associated to changes on gene expression and no changes were reported on *Dnmt3b* gene after chronic morphine treatment (**Supp Fig. 4**).

Transcriptomic analysis after chronic morphine treatment highlights a significant reduction in the expression of the *Dnmt1* maintenance methylase, as well as a significant increase in the expression of the *Tet1* demethylase gene. These RNA-seq results were validated by *RT-qPCR*, confirming the veracity of the RNA-sequencing results (**Fig. 3E**). In order to evaluate the impact of *Dnmt1* and *Tet1* gene expression alterations on DNA methylation pattern, global DNA methylation as well as hydroxymethylation levels were measured by mass spectrometry (LC-MS/MS) after chronic morphine treatment. Consistent with the gene expression results, chronic morphine treatment caused an overall decrease on DNA methylation together with an increase on hydroxymethylation in mESC (**Fig. 3F**). Specifically, morphine induced an hypomethylation on 5% of cytosines, meanwhile the overall level of hydroxymethylation underwent a significant increase of 15%. These results (LC-MS/MS) are in full agreement with those observed with WGBS technique, where chronic morphine treatment caused a global decrease in DNA methylation. Chronic morphine treatment, therefore, leads to a decrease in cellular methylation levels, generating a global hypomethylated state of the genome in mESC.

## DISCUSSION

Morphine is known to impact normal embryo development by affecting the neural tube, frontal cortex and spinal cord development, and, as a consequence, delaying nervous system development (Sahraei et al., 2013). Although morphine is an addictive drug able to cross the placental barrier and reach the embryo (Kazemi et al., 2011; Levitt, 1998), it is not fully understood how morphine specifically affects neurogenesis and other physiological aspects of embryonic development. Embryonic stem cells are widely used to study the impacts of environmental stimuli on developmental biology due to their ability to indefinitely self-renew and differentiate into any cell type (Dvash et al., 2006; Dvash & Benvenisty, 2004; Gepstein, 2002). Our results prove that DNA methylation may be an important epigenetic mechanism, key to understanding how morphine affects early embryonic development, as DNA methylation largely determines the direction of cell differentiation and the proper functioning of cells (Correa et al., 2020; Smith & Meissner, 2013). Consistent with changes induced by other addictive drugs such as cocaine (Novikova et al., 2008; Tian et al., 2012) and cannabinoids (Watson et al., 2015), WGBS techniques shown that chronic morphine treatment induces global DNA demethylation in mESCs, which might be mediated throw the activation of opioid receptors since they have been described in mESC (Dholakiya et al., 2016; Hahn et al., 2010). This global hypomethylation state is crucial for proper embryonic development, as it ensures the pluripotency of cells (Bansal et al., 2021). Morphine leads to a decrease in global DNA methylation that, together with a global downregulation of H3K27me3 (Muñoa-Hoyos et al., 2020), may be able to inactivate repressive microenvironments and promote active chromatin regions in mESCs. The hypomethylation induced by morphine can maintain a pluripotency state of the mESC and avoid cell differentiation, which may explain the retardation on nervous system development (Sahraei et al., 2013). Morphine impacts the DNA methylation distribution of key genes important for cell differentiation and development DNA conformation or metabolic processes beyond the regulation of their gene expression, so the role of DNA methylation is more complex and nuanced than has been thought.

DNA methylation might mediate the genomic response to morphine at genes involved in epigenetic regulation of gene expression, chromosome organization, cell cycle or metabolic process among others. Specifically, upregulation of *Tet1* gene may explain the global hypomethylation induced by morphine, as it is considering a demethylating enzyme (Wu & Zhang, 2014; Zhao & Chen, 2013). Consistent with other epigenetic marks such as H3K27me (Muñoa-Hoyos et al., 2020), morphine may have a self-regulation mechanism that modifies *Tet1* gene expression by modulating the methylation pattern of its promoter, thereby regulating its own expression through positive feedback. DNA demethylation is also a key mechanism in embryogenesis to maintain pluripotency (Montgomery et al., 2024). This process occurs through an active demethylation by TET enzymes, which leads to the oxidation of the methyl group (5mC) to hydroxymethylcytosine (5hmC) (Wu & Zhang, 2014; Zhao & Chen, 2013). Chronic morphine treatment can induce DNA hypomethylation through an active mechanism, as fact is also aligned with a significant rise in global DNA hydroxymethylation levels.

Besides active demethylation, passive mechanisms may also play an important role in overall morphine-induced demethylation (Chen & Riggs, 2011; He et al., 2017; Zhou et al., 2009). Our results prove an upregulation of *Dnmt3l* gene expression in response to morphine treatment, which is consistent with a downregulation of DNA methylation levels at gene promoter. DNMT3L, is required for germ line DNA methylation, although it is inactive as a DNA methyltransferase per se. Previous studies have shown that DNMT3L physically associates with the active de novo DNA methyltransferases, DNMT3A and DNMT3B, and stimulates their catalytic activity (Guenatri et al., 2013; Kareta et al., 2006). RNA-seq data confirm that morphine decreases *Dnmt1* gene expression but not *Dnmt3a nor Dnmt3b*. Due to the fact that the maintenance of DNA methylation patterns is predominantly carried out by DNMT1, while de novo methylation is primarily established by DNMT3A and DNMT3B methyltransferases (E. Li et al., 1992), morphine may compromise the maintenance of DNA methylation in mESCs, but its impact on *de novo* methylation may be limited. Although the sensitivity of DNA methylases to morphine is evident (Fan et al., 2019, Fan et al., 2021), it does not alter the promoter methylation status of these genes, suggesting that the decrease in *Dnmt1* gene expression can negatively affect the maintenance of methylation levels, leading to passive demethylation, but not through a DNA methylation mediated genomic response. Our findings confirm that morphine induces DNA demethylation through both active and passive mechanism, leading the already hypomethylated genome of mESC (Ito et al., 2011; Iyer et al., 2009; Pastor et al., 2013; Tahiliani et al., 2009; Wu & Zhang, 2014), to an even more hypomethylated state. The alteration of DNA methylation pattern, together with the inhibition of the PRC2 repressive machinery that leads to a global alteration in H3K27me3 chromatin organization, morphine promotes an aberrant transcriptome in mESCs, which may contribute to abnormal embryo development (Karanikolas et al., 2009; X. Li et al., 2009; Varambally et al., 2002).

To summarize, our results provide insights into how transcriptional changes induced by morphine may be mediated by DNA Methylation. Morphine disrupts selective target genes related to epigenetic regulation, chromosome organization, cell cycle and, particularly, to cell development and differentiation through alterations in DNA methylation pattern. Using both active and passive mechanisms that involve the action of demethylases and DNA methylases, chronic morphine treatment causes *in vitro* global hypomethylation in mESC, that can maintain the pluripotency state of the mESC. This morphine-induced DNA methylation changes might be key to the phenotypic changes that occur in adulthood. In fact, morphine-induced phenotypic changes in behavior, dopamine signaling pathways, synaptic plasticity, and neuronal structures (Byrnes et al., 2011; Chorbov et al., 2011; Sarkaki et al., 2008; Yohn et al., 2015), which have a strong potential to persist into adulthood (Sadraie et al., 2008; Saeidinezhad et al., 2021; Sahraei et al., 2013). Further experiments are needed to understand the mechanisms underlying morphine-induced heritable effects, which will be crucial for establishing the foundations of cellular memory in response to external stimuli during embryo development.

## METHODS

### Cell culture and treatment

Mouse embryonic stem cells (mESCs; Oct4-GFP cell line) (PCEMM08, PrimCells) were cultured with feeder-free medium on 0.1% gelatine-coated (Sigma) cell-culture dishes. The Knock Out Serum DMEM (Gibco) was supplemented with 15% KSR (Gibco), 1% sodium pyruvate (Sigma), 1% non-essential amino acids (Sigma), 1% penicillin– streptomycin (Sigma), 1% l-Glutamine (Sigma) and 0.07% β-mercaptoethanol (Sigma). The LIF+2i condition was obtained as follows: 1000 U/ml leukaemia inhibitory factor (LIF) (Sigma), 10 mM PD0325901 (Stemgent) and 30 mM CHIR99021 (StemCell). All experiments performed did not have the cells attached for more than 2 days. Therefore, the cells were passaged with 48-h intervals using trypsin (TrypLE Express Enzyme 1x, Thermo Fisher) for detachment. We were able to analyse the expression of GFP as an internal control of stemness, and thereby prevent differentiation. For chronic morphine treatment, mESCs were grown in their respective medium supplemented with %0.9 (p/v) NaCl and 10 μM morphine (Alcaliber) for 24h; control and treatment conditions, respectively. The cells were collected for further experimentation.

### Cell lysates, DNA extraction and quality measurement

To obtain mESC’s DNA, non-treated (control) and treated cells were lysated using DNA lysis Buffer (100 mM Tris-HCl, 5 mM EDTA, 200 mM NaCl and 0.2% SDS) supplemented with Proteinase K (AM2546, Thermo Fisher Scientific) at 100 mg/ml. Samples were incubated overnight under gentle shaking. The next day, the samples were treated with 5 μl RNAse at 10 mg/ml (R5125, Sigma) for 1-h at 37 °C. DNA extraction from all the samples was performed using a classic phenol-chloroform/isoamyl methodology with phenol (P4557, Sigma), chloroform (CL01981000, Scharlau) and isoamyl alcohol (BP1150, Fisher BioReagents). After the extraction, the DNA concentration and purity were evaluated by measuring the 260/280 absorbance ratio in a Nanodrop Spectrophotometer ND-1000 (Thermo Fisher Scientific).

### WGBS. DNA methylation analysis by Whole Genome Bisulfite Sequencing

The extracted DNA was sonicated to generate 300bp fragments (Soniprep 150). Those fragments, once denatured, they support a bisulfite treatment for later analysis. DNA bisulfite treatment was made with EZ DNA Methylation-Lightning Kit. DNA fragments were subjected to library construction using a KAPA Library Preparation Kit and xGenTM Methyl UDI-UMI Adapters. The library quality was confirmed using Agilent 2100 Bioanalyzer DNA 7500 assay 0. After library preparation, 4× multiple sequencing was performed (Paired End) on an Illumina NovaSeq 6000 S4 sequencer with a minimum of 50.000 reads for each of the two replicates per sample. The quality of the reads was verified using FASTQC analysis, and good overall alignment rate was confirmed for DNA methylation. Spearman correlation confirmed the reproducibility of the two biological replicates, and all genome cytosines in both duplicates of each sample were considered for further analyses.

### Bioinformatics analyses of WGBS’s data, and WGBS and RNA-sequencing base data integrative analyses

The whole bioinformatics process was done with the help of Bismarck and two different statistical programs. A FastQC High Throughput Sequence QC Report (v.0.11.6) was used to assess the quality of the FASTQ files from the WGBS, showing a quality score greater than 30. Library fragments adapters were cut with Trim Galore! (v.0.6.2), and consequently, reads obtained were concatenated with Cat (V.8.22). WGBS data was mapped to the UCSC mm10 reference genome using Bismarck (v.0.22.1), and the resultant binary equivalent files were sorted and indexed for quicker access using SAMtools (v.14.0). Spearman correlation analysis verified the reproducibility of each sample type. For the identification of *differentially methylated cytosines* (DMC) two different statistical tools were chosen: *edgeR* (v.3.32.1) and *methylkit* (v.1.16.1). The PCA (principal component analysis) in each of the tools confirmed the similarity between replicates and the differences between samples (morphine vs. control) (data not shown). After data normalization, the specific methylated cytosines that were identified by both statistical tools (*edgeR* and *methylkit*) coincided in 56%, a percentage that increase when we focus on analysing the genes that are affected (90% of genes). Considering the large amount of data, and in order to make a selection of the most significant results, we set different thresholds for each of the tools; minimum methylation-percentage difference of 25% (*methylkit*), and minimum fold-change difference of 2 (*edgeR*), and all these methylated cytosines differences have a p value of ≤ 0.05, considering differentially enriched. The used RNA-sequencing data was obtained previously through our investigation group with the same experimental procedure, and all these information is available (**RNA-seq - GEO storage: GSE151234**). WGBS and RNA-sequencing integrative analyses were performed using *Venny tools* (v.2.1.0), and *The Gene Ontology Resource* from the *GO Consortium* (v.16.1.0) (https://geneontology.org/) was used to identify the biological functions. The methylation and gene expression landscape are viewable via UCSC genome browser.

### LC-MS/MS. Quantification of DNA methylation by mass spectroscopy

The extracted DNA was enzymatically hydrolysed (DNA Degradase Plus; E2020, Zymo Research), and aliquoted samples (10 μl containing 50 ng of digested DNA) were run in a reversed-phase UPLC column (Eclipse C18 2.1 × 50 mm, 1.8 μm particle size, Agilent), equilibrated and eluted (100 μl/min) with water/methanol/formic acid (95/5/0.1, all by volume). The effluent from the column was added to an electrospray ion source (Agilent Jet Stream) connected to a triple quadrupole mass spectrometer (Agilent 6460/6400 QQQ). The machine was operated in positive ion multiple reaction monitoring mode using previously optimized conditions, and the intensity of specific MH+→fragment ion transitions was measured and recorded (5mCm/z 242.1 → 126.1, 5 hC 258.1 → 142.1 and dC m/z 228.1 → 112.1). The measured percentages of 5mC and 5hmC in each experimental sample were calculated based on the MRM peak area divided by the combined peak areas for 5mC plus 5hmC plus C (total cytosine pool).

### Real-time PCR (RT-qPCR)

The total mESCs RNA was extracted with Nucleozol reagent (Macherey-Nagel) according to the manufacturer’s instructions. Concentration was determined by measuring absorbance at 260 nm. Purity was assessed by the 260/280 nm absorbance ratio. All samples were reverse transcribed to DNA using the iScript cDNA synthesis Kit (Invitrogen). Real-time quantitative PCR analysis was performed using iTaq Universal SYBR Green SuperMix (Applied Biosystems, California USA) in PerkinElmer CFX96 Real Time Detection System (BioRad). The PCR profile was as follows: 39 cycles at 95 °C for 10 mins (denaturation), 20 secs at 95 °C (hybridization) and 1 min at 59 °C (annealing and extension). PCR amplification was repeated by more than three independent experiments in triplicates. The relative fold induction was quantified by the *ddCT* method. The most stable reference genes, glyceraldehyde 3-phosphate dehydrogenase (*Gapdh*) and pyruvate carboxylase (*Pcx)*, were used as housekeeping genes for data normalization. The used primer sequences are the following: *DNA methyltransferase 1* (*Dnmt1*), forward (F) 5’-GCC AGT TGT GTG ACT TGG AA-3′ and reverse (R) 5′-GTC TGC CAT TTC TGC TCT CC -3′; *DNA methyltransferase 3A* (*Dnmt3a*), forward (F) 5′-AAA CTT CGG GGC TTC TCC T-3′ and reverse (R) 5′-ATG GGC TGC TTG TTG TAG GT-3′; *DNA methyltransferase 3B* (*Dnmt3b*), forward (F) 5′-ACT TGG TGA TTG GTG GAA GC-3′ and reverse (R) 5′-CCA GAA GAA TGG ACG GTT GT-3′; *Tet methylcytosine dioxygenase 1* (*Tet1*), forward (F) 5’-TGC TCC AAA CTA CCC CTT ACA-3′ and reverse (R) 5′-CCC TCT TCA TTT CCA AGT CG-3′; *Tet methylcytosine dioxygenase 2* (*Tet2*), forward (F) 5′-TGG CTA CTG TCA TTG CTC CA-3′ and reverse (R) 5′-TGT TCT GCT GGT CTC TGT GG-3′; and *Tet methylcytosine dioxygenase 3* (*Tet3*), forward (F) 5′-CGC CGT GAT TGT TAT CTT GA-3′ and reverse (R) 5′-TGT TAG GGT CTT TGC CTT GG-3′; *Glyceraldehyde 3-phosphate dehydrogenase* (*Gapdh*), forward (F) 5′-TAT GAC TCC ACT CAC GGC AAA TT-3′ and reverse (R) 5′-TCG CTC CTG GAA GAT GGT GAT-3′; *Pyruvate Carboxylase* (*Pcx*), forward (F) 5′-CAA CAC CTA CGG CTT CCC TA-3′, reverse (R) and 5′-CCA CAA ACA ACG CTC CAT -3′.

## ACKNOWLEDGMENTS

The authors particularly acknowledge SGIker resources of UPV/EHU for technical support with the computational calculations, which were carried out in the Arina informatic cluster.

## AUTHOR’S CONTRIBUTION

MA, IMH, IUA and IC carried out the experiments and generated data. IMH performed the bioinformatic analysis. NS designed the study. NS and MA discussed the data and wrote the manuscript. The authors contributed to the discussion, read and approved the final manuscript.

## AVAILABILITY OF DATA AND MATERIAL

Authors consent to the availability of data and materials. The raw data have been deposited in NCBI Sequence Read Archive (SRA) through the Gene Expression Omnibus

