## Supplementary material for "Chronic morphine treatment leads to a global DNA hypomethylation via active and passive demethylation mechanisms in mESCs": Supplemetary information

#### Supplementary Figure 1

**A**

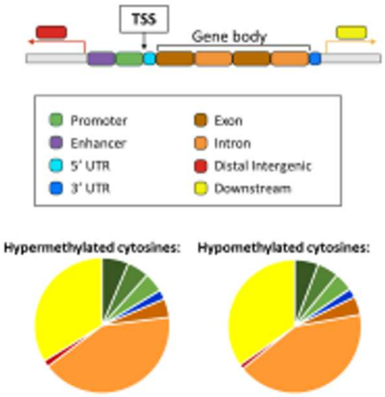


| | | DMC - Hypermethylations | DMC - Hypomethylations |
|---|---|---|---|
| Promoter – <=1kb | | 6.4 % | 5.8 % |
| Promoter – 1-2kb | | 5.2 % | 5.2 % |
| Promoter – 2-3kb | | 4.7 % | 4.5 % |
| 5' UTR | | 0.0 % | 0.1 % |
| 3' UTR | | 2.2 % | 2.2 % |
| Exon | | 4.4 % | 4.4 % |
| Intron | | 41.7 % | 42.2 % |
| Downstream | | 1.5 % | 1.1 % |
| Distal intergenic | | 33.8 % | 34.8 % |

**B**

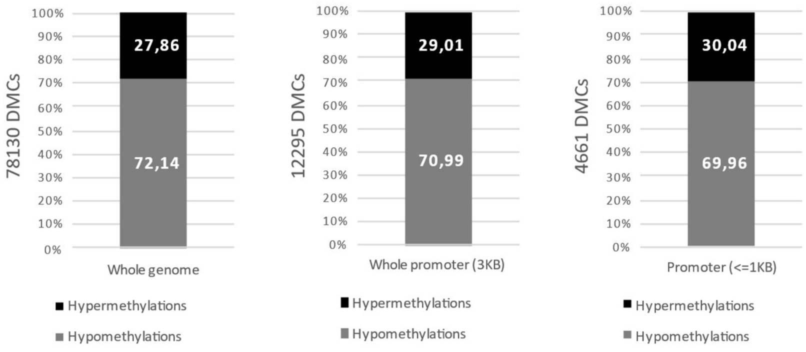


**Supplementary fig. 1. Analysis of the distribution of DMCs according genome features. (A)** Schematic image of the gene structure, and distribution of DMZ with respect to these gene areas, distinguishing hypermethylated and hypomethylated cytosines; **(B)** Graphs specifying percentage values of hypermethylations and hypomethylations as it approaches the promoter area: DMCs of the whole genome, DMCs that are located less than 3kb from the promoter, and DMCs of the promoter area.

Supplementary Figure 2

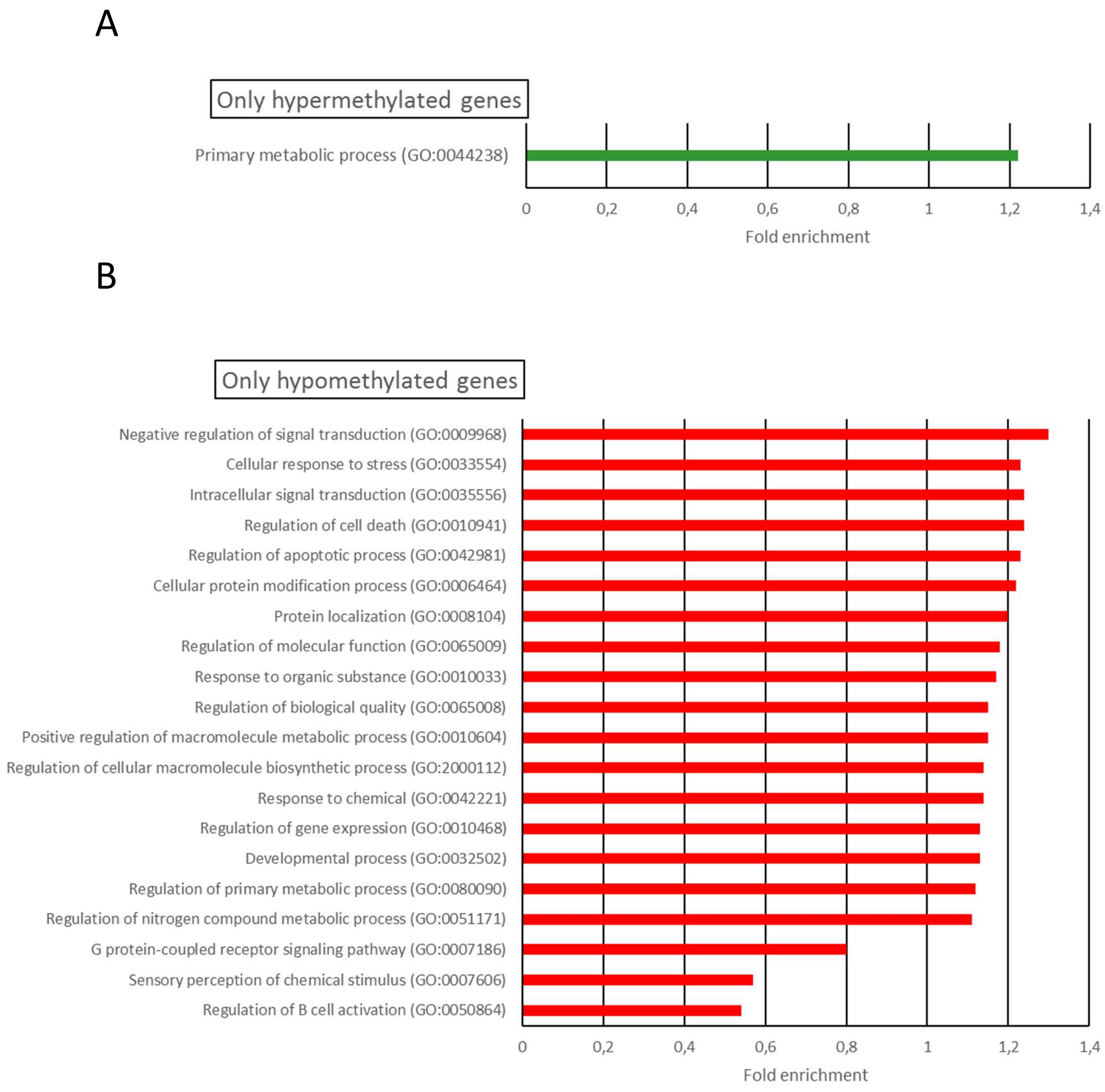


**Supplementary fig. 2. Functional enrichment analyses of DMC after chronic morphine treatment.** Gene Ontology analysis showing the top biological functions for **(A)** exclusively hypermethylated genes and **(B)** exclusively hypomethylated genes. All gene ontology analyses were corrected using Fisher's type test (0,05>p-value).

### Supplementary Figure 3

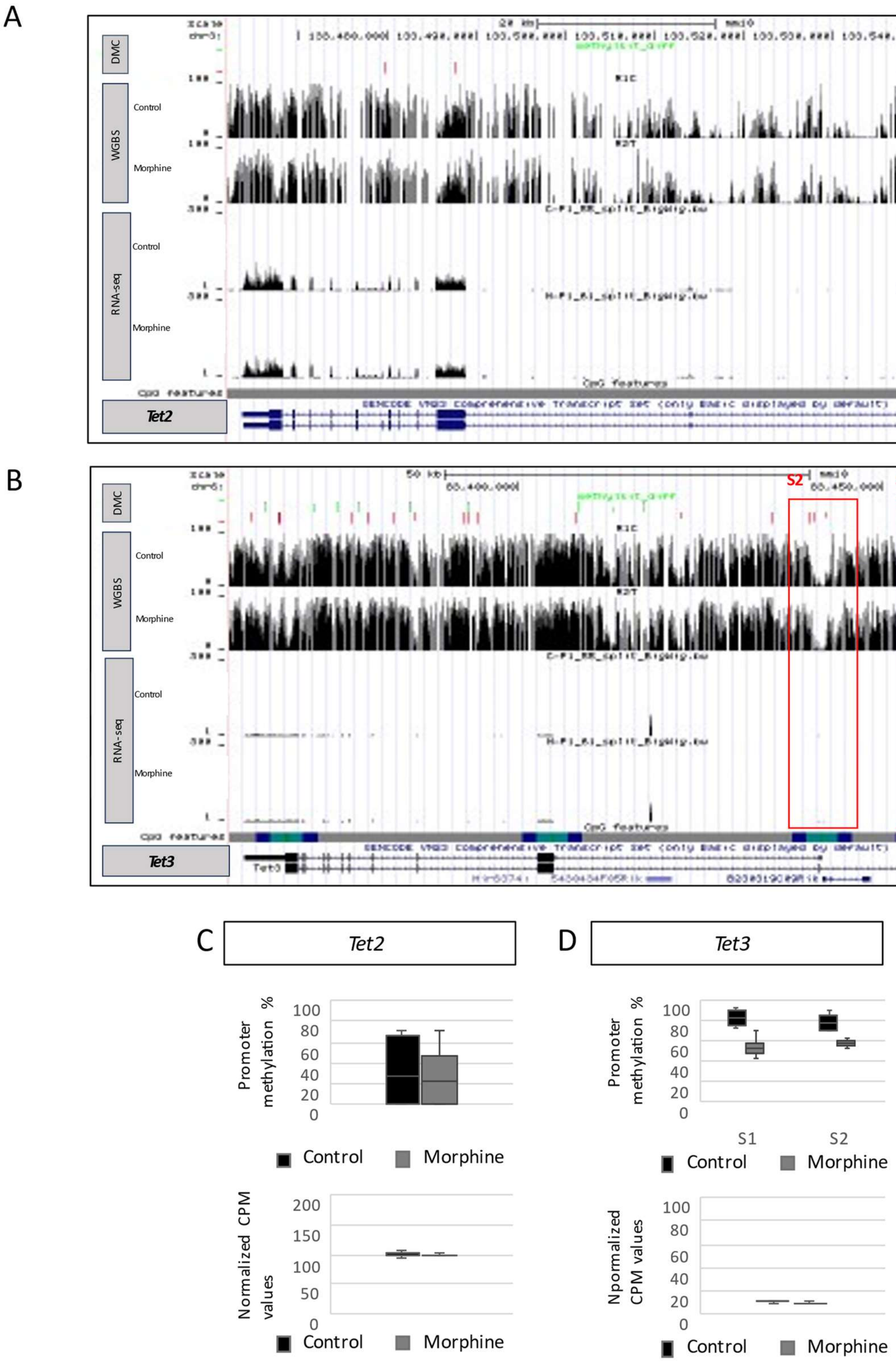

**Supplementary fig. 3. Effect of chronic morphine treatment on DNA demethylating proteins.** RNA-seq and WGBS track for **(A)** DNA demethylating enzyme *Tet2* gene and **(B)** *Tet3* gene. CpG features track was composed by CpG islands in green, shores in light blue, shelfs in dark blue and open sea in grey. Red boxes point out the enrichment and gene expression change at promoters. Box and whisker plot showing the percentage of methylation at promoters and CPM values for **(C)** *Tet2* gene and **(D)** *Tet3* gene after chronic morphine treatment.

### Supplementary Figure 4

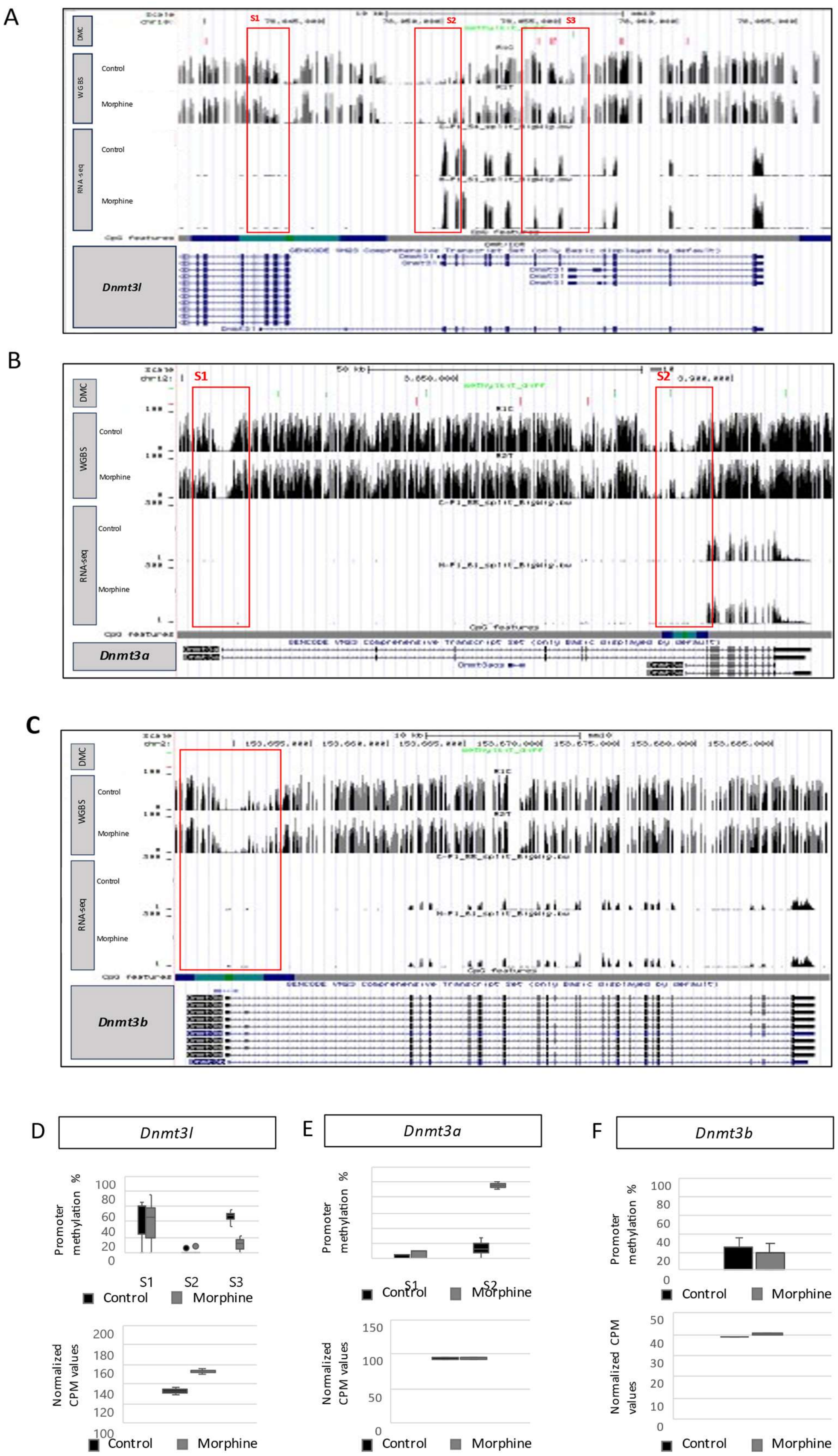

**Supplementary fig. 4. Effect of chronic morphine treatment on DNA methylation proteins.** RNA-seq and WGBS track for **(A)** DNA methyltransferase *Dnmt3l* gene, **(B)** *Dnmt3a* gene, and **(C)** *Dnmt3b* gene. CpG features track was composed by CpG islands in green, shores in light blue, shelfs in dark blue and open sea in grey. Red boxes point out the enrichment and gene expression change at promoters. Box and whisker plot showing the percentage of methylation at promoters and CPM values for **(D)** *Dnmt3l* gene, **(E)** *Dnmt3a gene* and **(F)** *Dnmt3b* gene after chronic morphine treatment.
